# Live tracking of inter-organ communication by endogenous exosomes in vivo

**DOI:** 10.1101/380311

**Authors:** Frederik J Verweij, Celine Revenu, Guillaume Arras, Florent Dingli, Damarys Loew, Gautier Follain, Guillaume Allio, Jacky G. Goetz, Philippe Herbomel, Filippo Del Bene, Graça Raposo, Guillaume van Niel

## Abstract

Extracellular vesicles (EVs) are released by most cell types but the definitive demonstration of their functional relevance remains challenging due to the lack of appropriate model organisms. Here we developed an *in vivo* model to study EV physiology by expressing CD63-pHluorin in zebrafish embryos. A combination of microscopy techniques and proteomic analysis allowed us to study the biogenesis, composition, transfer, uptake and fate of individual endogenous EVs *in vivo*. We identified an exosome population released in a syntenin-dependent manner from the Yolk Syncytial Layer into the blood circulation. These exosomes were specifically captured, endocytosed and degraded by patrolling macrophages and endothelial cells in the Caudal Vein Plexus (CVP) in a scavenger receptor and dynamin-dependent manner. Interference with exosome secretion affected CVP growth, supporting their trophic role. Altogether, our work provides a unique model to track in vivo inter-organ communication by endogenous exosomes at individual vesicle level and high spatio-temporal accuracy.

**Highlights:**

- Single endogenous EVs can be live-visualized in the whole embryo with CD63-pHluorin
- In the YSL, syntenin regulates exosome release into the blood for their propagation
- YSL exosomes reach the tail to be taken up by macrophages and endothelial cells
- Uptake is scavenger receptor and dynamin-dependent and provides trophic support

**Blurb:** We propose zebrafish embryos expressing a fluorescent reporter for exosomes as a relevant model organism to live-track production, journey and fate of individual extracellular vesicles in vivo. Our model allows investigation of the composition of EVs and the molecular mechanisms controlling their biogenesis and fate and functions in receiving cells.

## MAIN TEXT

Extracellular vesicles (EVs) form a collection of vesicles that include exosomes, 50-150nm diameter membranous vesicles of endosomal origin, and microvesicles (MVs), 50-500nm diameter membranous vesicles that bud from the plasma membrane (PM). Exosomes and MVs are released by a wide variety of cell types and are found in all organism investigated so far (van Niel et al., 2018). Their implication in an increasing number of major physiological and pathological processes supports their putative role as essential intercellular messengers (Tkach and Théry, 2016). EVs are notably involved in embryonic development, angiogenesis, immune regulation, and trophic support under physiological conditions. Moreover, in pathological contexts they have been shown to support metastasis and the development of metabolic, cardiovascular and neurodegenerative diseases among others (Bian et al., 2014; Bobrie et al., 2011; Costa-Silva et al., 2015). Most of the data on the function of EVs and particularly exosomes, however, is gathered from transformed cancer cells and relies on the use of a heterogeneous population of exosomes purified from cell culture supernatant or liquid biopsies. Consequently, the propagation of endogenous exosomes in vivo are largely unknown and data on their biogenesis and role in normal developing tissue and adult tissue homeostasis distinctly lags behind. The understanding of EV biogenesis, transfer and fate in recipient cells is, however, of prime importance not only from a fundamental point of view, but also to assess their relevance in pathological conditions as well as for their use as early biomarkers of pathologies or as therapeutic delivery systems (Fais et al., 2016). This apparent hiatus is largely due to the shortage of suitable models to visualize and track single endogenous EVs, and in particular exosomes *in vivo* from their site of production to their final destination in target cells (Hyenne and Goetz, 2017; Tkach and Théry, 2016). Previous studies visualizing EV release and transfer have been limited to the detection of larger extracellular structures (Lai et al., 2015; Zomer et al., 2015) or their transfer from transplanted transformed cells (Zomer et al., 2015).

Our main aim was to develop a unified model to study single EV release, transfer and function, which combines *in vivo* imaging with sub-cellular resolution while retaining the *in toto* imaging scale. Due to their small size and transparency, zebrafish embryos elegantly match the single-cell precision of C. elegans research in a vertebrate system with relevant architectural homology with mice and human (Megason, 2009). We show here that a zebrafish cell line secretes EVs containing classical exosome markers such as CD63, thus allowing the study of EV biogenesis and physiology in the context of the whole organism with the CD63-pHluorin reporter in zebrafish embryos. This fluorescent reporter is specifically targeted to (late-) endosomes and secreted on exosomes, allowing live visualisation of exosome release in single cells *in vitro* (Verweij et al., 2018). Ubiquitous expression of CD63-pHluorin in zebrafish embryos revealed massive release of EVs *in vivo* that could be tracked in the blood flow of the whole animal up to their final destination. Specific expression of CD63-pHluorin in the Yolk Syncytial Layer (YSL) revealed a major source of EVs. The YSL is a multinucleate cell layer enveloping the yolk that plays crucial roles in embryonic patterning and morphogenesis at early development stages and has essential metabolic and nutrient transport functions during embryonic and larval stages (Carvalho and Heisenberg, 2010). We could track the YSL-derived EVs, further identified as exosomes, once released in the blood flow. Using a combination of live embryo imaging and electron microscopy, we show that YSL derived exosomes can go through the whole organism and are specifically arrested by macrophages and endothelial cells of the caudal vein plexus. Both cell types take up YSL derived exosomes in a dynamin-dependent manner for degradation in lysosomes. Endocytosis occurred through interaction with Class-A Scavenger Receptors of endothelial cells. Interfering with the syntenin-a-dependent biogenesis of exosomes affected vasculogenesis of target cells, supporting a physiological trophic role of YSL derived exosomes. Altogether, these data reveal for the first time the secretion, journey and the target of individual endogenous exosomes secreted *in vivo*.

## RESULTS

### Zebrafish cells secrete EVs containing classical exosome markers

To investigate whether zebrafish could serve as a relevant model to study exosomes, we first assessed the presence of EVs in zebrafish embryos 3 days post fertilization (3dpf) by electron microscopy (EM). EM analysis revealed the presence of 100-200nm diameter vesicles clustered and enwrapped or free in the extracellular space (**Fig 1A**). We also processed the cell-culture supernatant of the zebrafish fibroblast AB.9 cell line with a differential ultra-centrifugation protocol (Théry et al., 2006). By electron microscopy we detected various vesicles in the size range of exosomes (~100nm) (**Fig 1B**). Proteomic analysis on the same preparations identified a variety of zebrafish homologues of exosomal markers traditionally found in mammalian exosomes, including tetraspanins CD63, CD9, CD82, and several heat-shock proteins (HSPs), including HSP70 and HSP90 (**Fig 1C, Suppl. table I**) and similar to the markers identified in exosomes derived from Zmel cells, a zebrafish melanoma cell line (Hyenne *et al*., co-submitted). This demonstrates that zebrafish cells release EVs that are enriched in the CD63 zebrafish ortholog, and supports the application in zebrafish of a fluorescent reporter that we recently used to visualize MVB-PM fusion *in vitro*, hCD63-pHluorin (Verweij et al., 2018). Due to its pH-sensitivity, CD63-pHluorin primarily reports its ‘extracellular pool’ (i.e. PM and EVs) but obscures CD63 large intracellular pool present in acidic endosomal compartments (Sung et al., 2015), which facilitates the detection of smaller extracellular CD63 positive (CD63+) objects such as EVs. Injection of ubi:hCD63-pHluorin plasmid-DNA (pDNA) at the single-cell stage of Casper zebrafish embryos resulted in mosaic, transient expression of CD63-pHluorin in multiple cell types, including epithelial, endothelial, neuronal, red-blood- and muscle cells (**Fig 1D-E**). CD63-pHluorin associated fluorescence was localized at the PM of these cell types and to dotty structures potentially corresponding to CD63-pHluorin positive released EVs (**Fig 1E, arrows**). An additional benefit of using CD63-pHluorin, as previously shown in cell lines (Verweij et al., 2018) is the possibility to visualize and estimate the frequency of MVB-PM fusion events *in vivo*. Current technical limitation of imaging limited the detection of MVB-PM fusion events *in vivo* in flat and stretched cells close to the dermis such as long-stretched fibroblast-like cell type (**Fig 1F**). Overall, we detected a low number of fusion events in vitro (~1 every 10 minutes) with a duration comparable to our observation in various cell lines *in vitro* (i.e. 20-30 sec) (Verweij et al., 2018).

**Figure 1.**
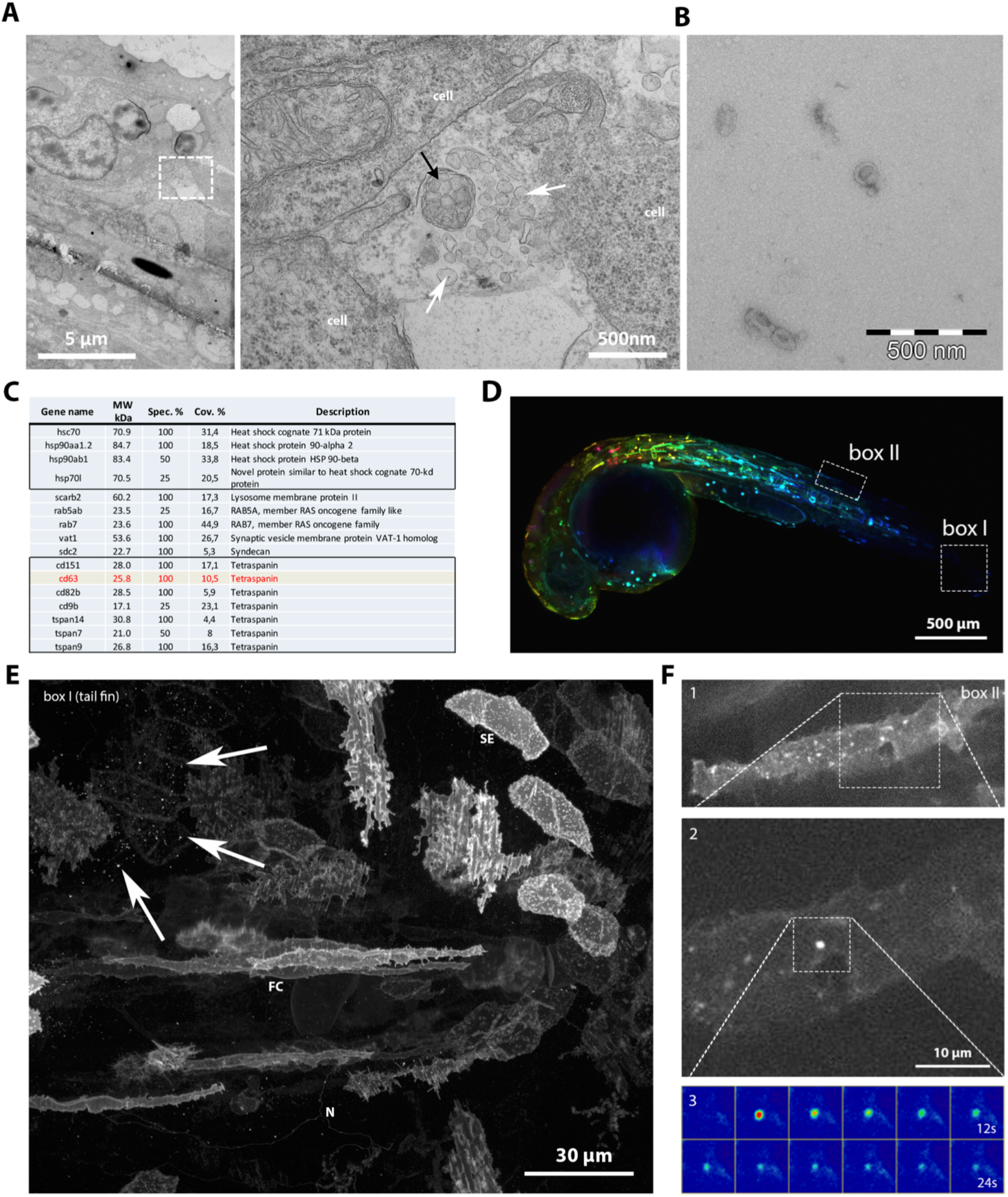
Description of the model. (A) Electron microscopic analysis of a 3dpf zebrafish embryo showing heterogeneous extracellular vesicles. White arrows indicate exosome-like vesicles, black arrow indicates an enwrapped cluster of vesicles. (B) Electron microscopy observation of exosomes purified by ultra-centrifugation from the supernatant of AB.9 fibroblast cells. (C) Mass-spec analysis of AB.9-derived exosomes (excerpt). (D) Transient mosaic expression of ubi:CD63-pHluorin in 35hpf zebrafish embryo, pseudo-coloured for depth. (E) Example of cell types in caudal fin targeted by transient mosaic expression of CD63-pHluorin (3dpf). Identified can be, among others, skin epithelial (SE), neuronal (N), fibroblast cells (FC). Arrows point at labelled EV. (F) Example of a fusion event observed in a fibroblast-like cell (1), with zoom in (2) and time-lapse heat map of the event (3).

### A major pool of CD63 positive EVs is present in the bloodstream of zebrafish embryos

One of the most commonly postulated assets of EVs is their potential as extracellular carrier long-range communication *in vivo* (Zomer et al., 2015) by entering the circulation (Tkach and Théry, 2016). In zebrafish embryos, the cardio-vascular system develops as early as at 24 hours post fertilization with a heart-beat and a first transient wave of erythrocytes (Hermkens et al., 2015). Strikingly, in 3dpf zebrafish larvae expressing CD63-pHluorin, we could detect a large pool of CD63-pHluorin positive structures carried by the blood flow in different parts of the vasculature throughout the body, including the brain and the gills (**Fig 2A-B; Suppl. Videos 1-3**). EM analysis on blood vessels of ubi:CD63-pHluorin plasmid (p)DNA expressing embryos confirmed the presence of numerous single EVs in the size range of exosomes (~100nm), but also a minor fraction of trapped vesicles and multivesicular structures that labelled positive for aGFP immune-gold (**Fig 2E-D**). While the initial observations were done on 3dpf embryos, we could detect CD63 positive EVs as early as 24h (1dpf), at the onset of the blood flow (data not shown). This observation opened the opportunity to track individual endogenous EVs in vivo.

**Figure 2.**
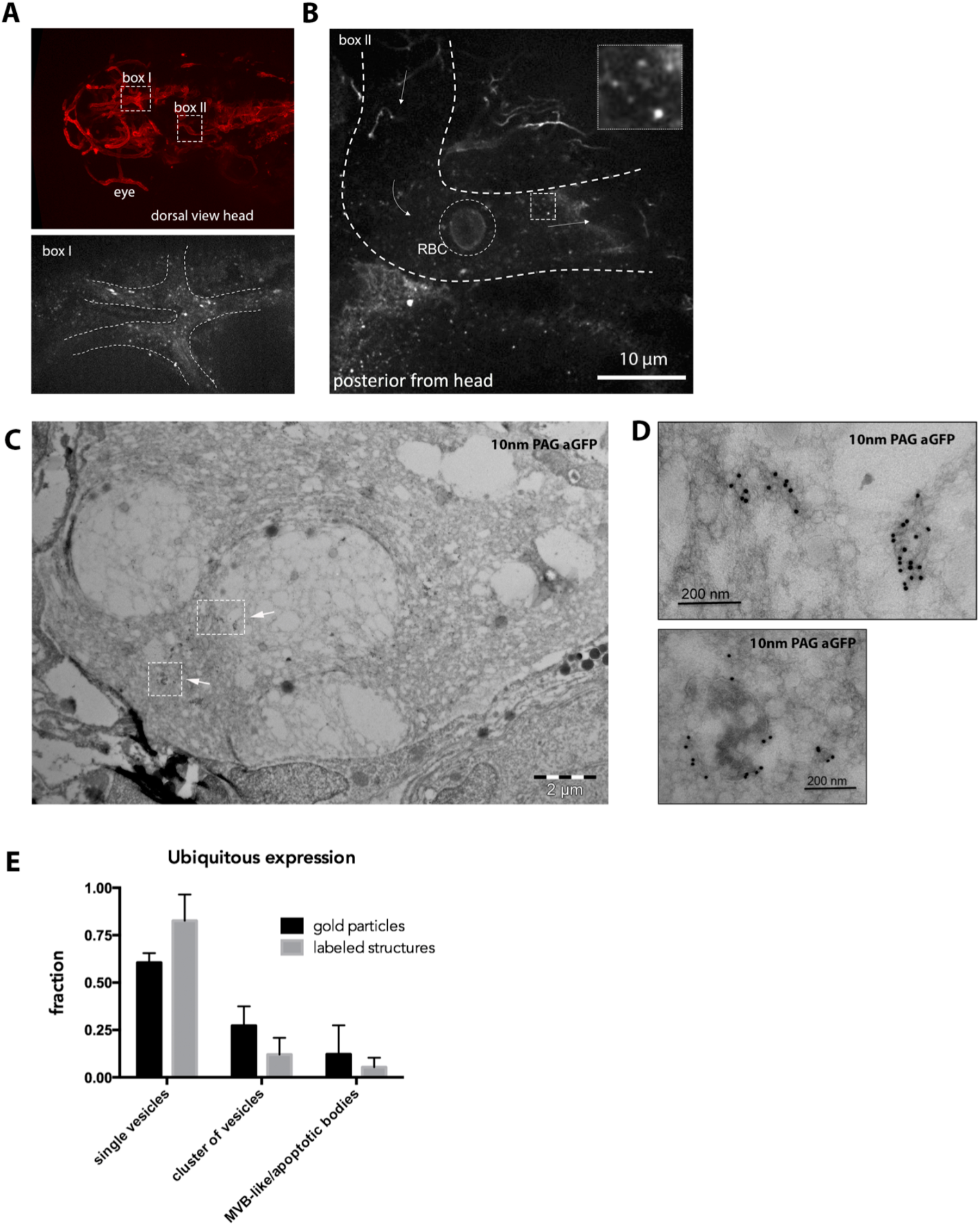
Exosomes in the blood flow. (A) Dorsal view of the vasculature in the head of a *Tg(kdrl:Hsa.HRAS-mCherry)* 3dpf zebrafish larva expressing CD63-pHluorin after injection of ubi:CD63-pHluorin pDNA at the 1-cell stage. Boxes I and II indicate areas of interest imaged for CD63-pHluorin, shown in a (lower panel) and b. (B) Still of time-lapse sequence shown in Sup Video 1, indicating the approximate vasculature wall (dashed white lines), a red blood cell (RBC), and in the dotted box a sample of extracellular vesicles seen in the bloodstream. Arrows indicate the direction of the blood flow. (C) EM on the vasculature of ubi:CD63-pHluorin pDNA injected fish (transient mosaic expression), labelled with gold particles directed to GFP (10 nm), showing (D) single-(upper panel) and clusters (lower panel) of vesicles, corresponding to the dotted boxes in c. (E) Quantification of vesicle appearance (single/clustered) of EVs as in (c).

### The yolk syncytial layer (YSL) releases CD63-pHluorin positive exosomes in the bloodstream

Next, we aimed to identify the cellular origin of the CD63 positive EVs detected in the bloodstream. We reasoned that, in view of the high number of EVs in the blood, the cell type/tissue of origin is likely in direct contact with the blood. The large surface area of the yolk syncytial layer (YSL) and their highly differentiated secretory apparatus, high lysosomal activity, and cytoplasmic multivesicular structures by electron microscopy (Walzer and Schönenberger, 1979) made a plausible candidate for the vast number of EVs detected in the blood flow. Video-enhanced Nomarski microscopy revealed numerous EVs in the large interstitial space between the YSL and the overlying epidermis, at and even before the onset of blood circulation (**Supp Fig 1A**). Injection of ubi:CD63-pHluorin pDNA directly in the YSL at the 1000-cell stage (4hpf) (**Supp Fig 1B**) allowed for tissue-specific expression of CD63-pHluorin and circumvents the need for tissue specific promoters (**Fig 3A**). Immuno-gold labelling for GFP on thin cryo-sections of YSL CD63-pHluorin expressing 3dpf zebrafish embryos showed a massive number of vesicles ranging from 80-200nm in size immediately above the YSL and in direct contact with the blood (**Fig 3B-C and Supp Fig 1C**). Consistently, CD63-pHluorin positive EVs were detected in the blood flow after site specific expression of CD63-pHluorin in the YSL (**Suppl. Video 4**), and EM analysis of the blood vessels confirmed the nature of the fluorescent structures as single vesicles enriched in CD63-pHluorin in the size range of exosomes (~100nm) (**Fig 3D**). Moreover, YSL derived EVs that were observed in the blood flow accumulated in the caudal vein plexus (CVP) only in embryos with a detectable bloodstream (**Fig 3E**). This indicates that YSL-derived EVs enriched in CD63-pHluorin are continuously released in the blood flow and can circulate through the entire embryo to reach their remote destination.

**Figure 3.**
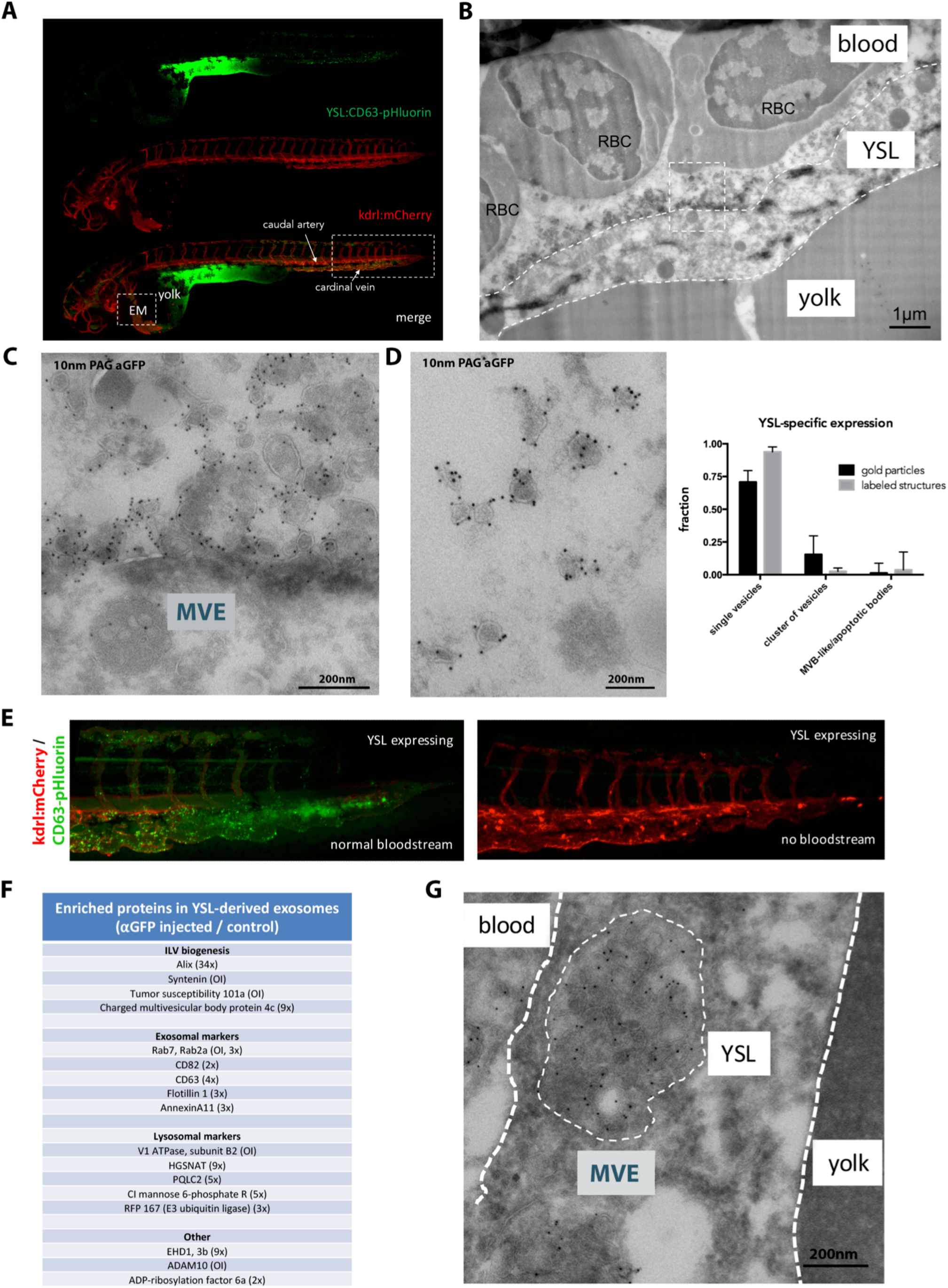
Origin and distribution of exosomes in the blood flow. (A) Fluorescence image of a 3dpf *Tg(kdrl:Hsa.HRAS-mCherry)* embryo injected with ubi:CD63-pHluorin in the YSL, green channel (top), red channel (middle) and merge (bottom). Box [EM] indicates the approximate region for EM analysis (shown in c). Box to the right indicates the area shown in e. (B) EM image of the YSL of a 3dpf *Tg(kdrl:Hsa.HRAS-mCherry)* embryo injected with ubi:CD63-pHluorin in the YSL labelled with gold particles directed to GFP (10 nm). Dashed white lines indicate the approximate membranes of the YSL. Dashed line box indicates area shown enlarged in d. RBC = red blood cell. (C) Zoom in of area indicated in (c), showing MVEs in the YSL, and vesicles labelling positive for gold particles directed to GFP (10 nm) on top of the YSL. (D) EM image of exosome-sized EVs found in the blood, labelled with gold particles directed to GFP (10 nm). Graph on the right: quantification of vesicle appearance as in Fig 2e (E) Zoom in on area indicated with rectangular box on the right in a. To the left, fish with blood flow, to the right, fish without blood flow, both expressing CD63-pHluorin in the YSL. (F) Mass-spec analysis of enriched proteins in YSL-derived EVs (excerpt). (G) EM image of an MVB in the YSL area, labelled with gold particles directed to GFP (10 nm).

Next, we characterized the nature of these fluorescent EVs by dissociating 3dpf CD63-pHluorin YSL expressing and non-expressing control embryos using Collagenase I treatment, and isolating EVs from the supernatant with a differential ultra-centrifugation protocol (Théry et al., 2006). Nanoparticle tracking analysis (**Supp Fig 1D**) of the 100 000g pellet of both conditions did not show significant differences in terms of size and number, suggesting that the expression of CD63-pHluorin in the YSL did not profoundly affect the release of EVs compared to basal condition. Immunogold labelling followed by electron microscopy observation of the CD63-pHluorin YSL expressing embryos derived pellet (**Supp Fig 1E**) confirmed the presence of vesicles heavily labelled for CD63-pHluorin with a morphology and a size (80-120nm) comparable to those observed by EM in the blood flow (**Fig 3D**). Both pellets were then further processed by immunoprecipitation against GFP using GFP-Trap agarose beads, followed by quantitative label-free proteomics analysis of the pull-down fraction. We quantified enrichment of specific proteins using non-injected zebrafish derived EV pellet as background. The most enriched proteins were syntenin-a, TSG101, Rab7, ADAM10 and V1 ATPase subunit B2 that were only detected in the injected YSL EV-fraction and Alix (34x). Other YSL EV-enriched proteins included CMHP4c, EHD1, Cation-Independent Mannose 6-phosphate Receptor, Flotillin 1, CD63, CD82, the lysosomal acetyltransferase HGSNAT and the lysosomal amino acid transporter PQLC2. (**Fig 3F, Supp Table II**). Of note, several of the enriched proteins are implicated in one of the machineries involved in exosome biogenesis in endosomes in vitro (Baietti et al., 2012). These data, strongly supporting the endosomal origin of the YSL-derived EVs, were further strengthened by the presence of large MVB structures containing CD63-pHluorin positive ILVs (**Fig 3G**) in the YSL, indicating that at least part of the YSL-EVs present all the characteristics of exosomes as previously defined (Gould and Raposo, 2013). Finally, these YSL-exosomes were enriched for Molecular function GO terms “Anion binding”, “Small molecule binding” and ‘Transporter activity’, the latter being characterized by the presence of several Solute Carrier Family proteins, including zinc and amino acid transporters slc30a1a and slc3a2b. YSL derived exosomes displayed similarities with the composition of the AB.9 caudal fibroblast derived exosomes and Zmel zebrafish cell line derived EVs (**Supp Table I**; (Hyenne *et al*., co-submitted)), as they both contain major actors of exosome biogenesis (Alix, TSG101, Chmp4C, tetraspanins). Nevertheless, physiological and pathological exosomes differed by their composition that could be linked to their respective (patho)physiology (**Supp Table II**). Altogether, these data strongly suggest that expression of CD63-pHluorin in the YSL reveals the release of labeled exosomes in the bloodstream of developing embryos.

### YSL-derived CD63 positive exosomes adhere to endothelial cells of the caudal plexus and are actively endocytosed by patrolling macrophages

Having determined a cell layer of origin and transportation route of CD63 positive exosomes, we live-tracked them in the blood vasculature to identify potential target cells. Our data revealed an accumulation of dotty signals (**Fig 3A**) likely corresponding to immobilized exosomes in the blood, in particular in the caudal part of the vasculature. This area comprises the caudal artery, and a complex, well-perfused, venous vascular network in between and originating from the cardinal vein, called the caudal vein plexus (CVP) (Isogai et al., 2001).

A maximum projection time-lapse movie of the VP at higher magnification suggested an interaction between the labelled exosomes and the vessel wall (**Fig 4A, Suppl Movie 5**). Indeed, high-speed imaging of the cardinal vein confirmed the transient interaction of exosomes passing through the blood vessels with the plasma membrane of endothelial cells (**Fig 4B, Suppl Movie 6**). To visualize the extent and the exact localization of exosomes that adhere in the caudal part of the vasculature, we created a heat-map overlay of 6 individual YSL CD63-pHluorin pDNA injected embryos 3dpf (**Fig 4C**). Strikingly, the vast majority of exosomes was arrested in the venous plexus and the caudal vein, but not the caudal artery. This behaviour is similar to what was observed for intravenously injected Zmel1 EVs (Hyenne et al., co-submitted), suggesting a common behavior for EVs in the blood flow of zebrafish embryos. In our high-magnification images we identified motile cells that crawled on the surface of the endothelial vessels and that contained labelled exosomes in intracellular compartments, likely resulting from their endocytosis (**Fig 4D**). Apart from endothelial cells, the lumen of the caudal vein plexus is home to a number of embryonic macrophages that continuously scavenge their environment and stand out by their relative high motility (Murayama et al., 2006). In *Tg(kdrl:HRAS-mCherry)* zebrafish embryos, these macrophages can generally be distinguished by their mCherry-positive compartments, probably as a result of endocytosis of (apoptotic) fragments of the mCherry-expressing endothelial cells as observed in motile cells accumulating labelled exosomes (**Fig 4D**). Identification of the motile cells internalizing labelled exosomes as patrolling macrophages was confirmed in a *Tg(mpeg1:mCherryF)* fish line, that specifically labels macrophages in red (**Fig4E, Supp Movie 7**). Subsequent time-lapse imaging of macrophages in the caudal plexus of *Tg(kdrl:HRAS-mCherry)* fish expressing CD63-pHluorin in the YSL revealed an active form of uptake of exosomes by macrophages that projected dendrites or pseudopodia able to interact and ‘harvest’ exosomes from the surface of endothelial cells (**Fig 4F; Supp Movie 8**). Similar ‘harvesting’ by patrolling macrophages was observed in Hyenne for melanoma-derived exogenous EVs (Hyenne *et al*., co-submitted), suggesting common mechanisms for uptake of EVs of distinct origin.

**Figure 4.**
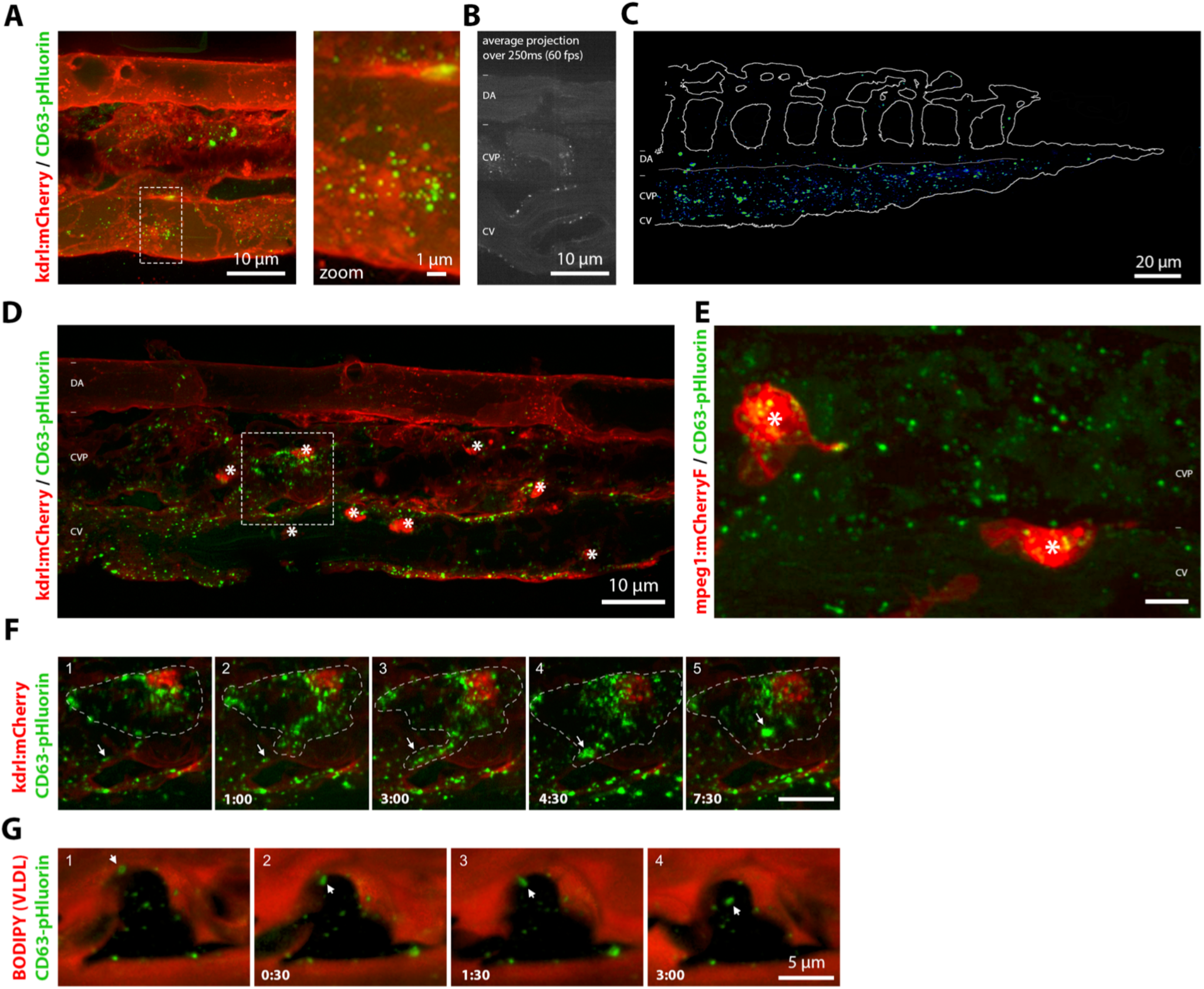
Distribution and target cells. (A) Still of time-lapse shown in Sup Video 5 of the CVP of a 3dpf *Tg(kdrl:Hsa.HRAS-mCherry)* embryo expressing CD63-pHluorin in the YSL. Dashed box indicates area shown at higher magnification to the right. (B) Average projection of 15 sequential frames of Sup Movie 6, showing dorsal artery (DA), CVP and the cardinal vain (CV) in 3dpf zebrafish embryos expressing CD63-pHluorin in the YSL. (C) Heat-map of CD63-pHluorin signal in CVP of 6 3dpf *Tg(kdrl:Hsa.HRAS-mCherry)* zebrafish embryos expressing CD63-pHluorin in the YSL. (D) CVP of a 3dpf *Tg(kdrl:Hsa.HRAS-mCherry)* embryo expressing CD63-pHluorin in the YSL. Stars indicate macrophages. Dashed box indicates area shown in Sup Movie 8 and in (f). (E) Still of Sup Movie 7, showing the CVP of a 3dpf *Tg(mpeg1:mCherryF)* expressing CD63-pHluorin in the YSL. Stars indicate macrophages. (F) Time-lapse of a macrophage in a 3dpf *Tg(kdrl:Hsa.HRAS-mCherry)* zebrafish embryo expressing CD63-pHluorin in the YSL, shown in Sup Movie 8. (G) Time-lapse of a BODIPY-C12 injected macrophage in a 3dpf zebrafish embryo expressing CD63-pHluorin in the YSL, shown in Sup Movie 8. Asterisks indicate macrophages.

To further investigate the specificity of the uptake of labelled exosomes by macrophages of the CVP, we injected the yolk of CD63-pHluorin YSL expressing Casper embryos at 1dpf with 1.5 ng/nl BODIPY-C12 (Miyares et al., 2014) to specifically label the high amount of VLDL particles produced by the YSL and released in the blood flow, and to visualize the lumen of the blood vessels (Anderson et al., 2011; Miyares et al., 2014). The BODIPY fluorescence was readily visible in the vasculature but was not overlapping with CD63-pHluorin labelling in the blood flow, confirming the nature of CD63-pHluorin positive EV as exosomes and not as derivatives of VLDL. Moreover, we did not observe any BODIPY signal in macrophages, while exosomes were again actively endocytosed (**Fig 4G; suppl Movie 9**). We did not observe an early decrease in fluorescence signal of CD63-pHluorin early (~3 minutes) after uptake (**Fig 4G**) while the first co-localization of melanoma derived EVs with Lysotracker appeared 10 minutes after their injection, suggesting targeting to acidic endosomal compartments. All in all, these data revealed a specific uptake of exosomes by macrophages in intracellular compartments that become acidic at later time point and/or that initially maintain connection with the cellular exterior.

### YSL-derived exosomes are taken up and degraded by endothelial cells

The specific interaction of labelled exosomes with the venous plexus and the posterior cardinal vein (**Supp Movie 5**) but not the caudal artery (**Fig 4C**), suggested potential implication of different shear forces exerted on EVs (Hyenne *et al*., co-submitted) between arterial and venous blood flows. Drug treatment decreasing pacemaker activity (Vermot et al., 2009), and consequently the shear force of the blood flow, did not lead to a visible increase of the association of exosomes with the caudal artery (**Sup Fig1F**). This indicates the implication of additional factors, such as expression of adhesion molecules, in this interaction.

We further analysed the dynamic of interaction between exosomes and the endothelium by using high-speed imaging of the caudal vein of YSL CD63-pHluorin pDNA injected embryos 3dpf. We observed rolling, arrest and accumulation of exosomes on the endothelial wall of the vein (**Fig 5A, Sup Movie 10**). Quantification of the duration of the interaction between exosomes and the venous vessel wall indicated that exosomes stay attached for a duration between 1 up to 50 minutes, with an average of 3 min (**Fig 5B**). This behaviour is similar to that of exogenous melanoma-derived EVs (Hyenne *et al*., cosubmitted) and reinforced potential molecular interactions between exosomes and specific receptors at the surface of venous endothelial cells.

**Figure 5.**
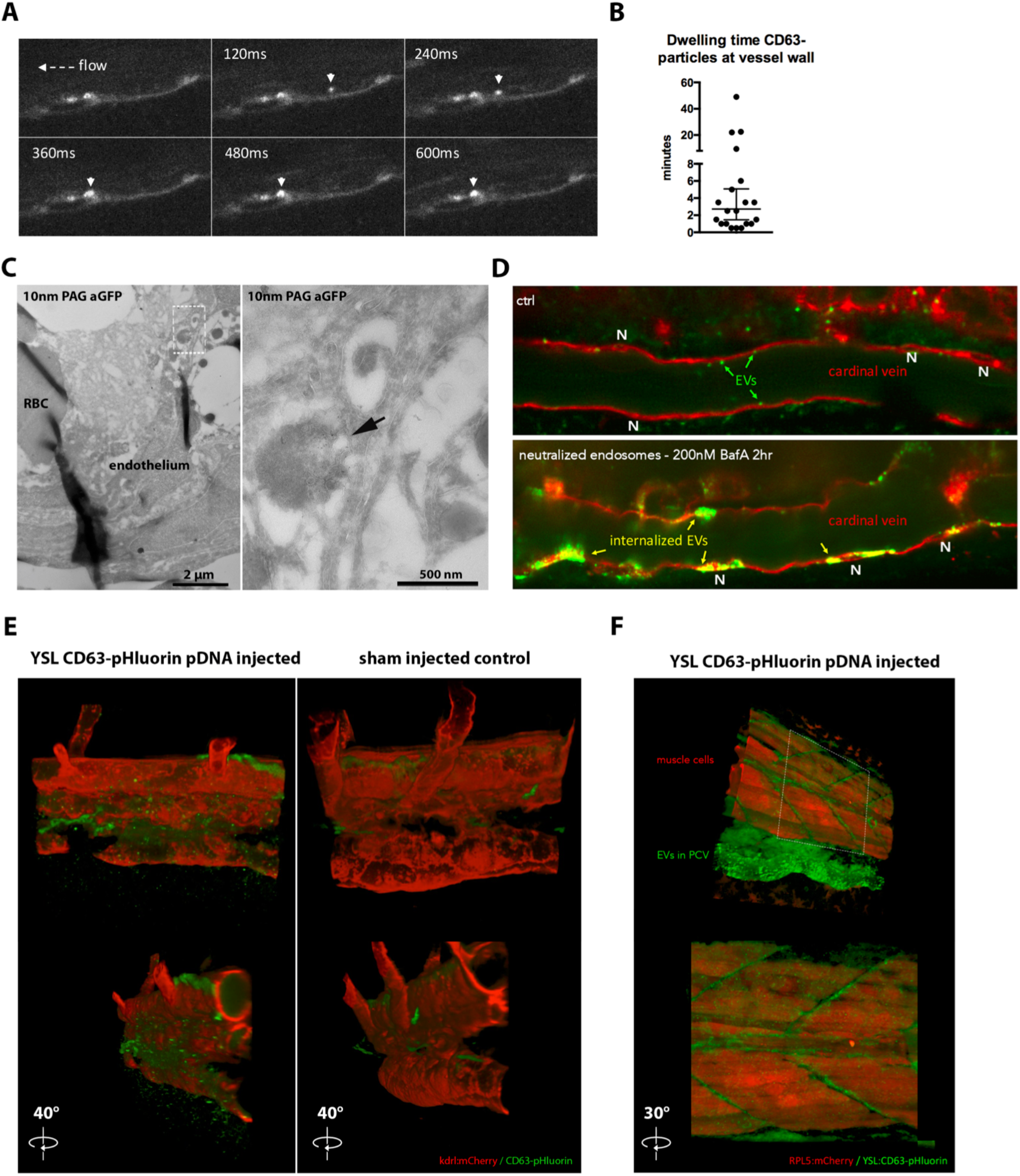
Interaction of YSL-EVs with endothelium and presence in ISF. (A) Time-lapse of Sup Video 10, showing the rolling and arrest of EVs in the cardinal vein of 3dpf zebrafish embryo expressing CD63-pHluorin in the YSL. (B) Quantification of dwelling time of CD63-pHluorin vesicles at the vascular wall. (C) EM image of the CVP of a 3dpf zebrafish embryo expressing CD63-pHluorin in the YSL, labelled with gold particles directed to GFP (10 nm). Insert indicates area shown at higher magnification to the right. Arrow indicates internalized CD63-pHluorin EVs. (D) Close-up of the cardinal vein of 3dpf *Tg(kdrl:Hsa.HRAS-mCherry)* zebrafish embryos expressing CD63-pHluorin in the YSL, control treated (DMSO) or treated with BafilomcinA (BafA) to neutralize acidic compartments. (E) CVP area of 3dpf *Tg(kdrl:Hsa.HRAS-mCherry)* zebrafish embryos injected with CD63-pHluorin pDNA or sham-injected in the YSL. (F) Mid-trunk segment of of 3dpf zebrafish embryos expressing CD63-pHluorin in the YSL and mCherry under the ubiquitous RPL5 promoter. At this exposure level in the red channel, mostly muscle cells are visible.

We observed that exosomes interacted transiently with endothelial cells and then disappeared (**Fig5B, Supp Video 5**), suggesting either their uptake by endothelial cells and targeting to acidic endosomal compartments, their re-entry into the blood flow or their dispersion outside of the vasculature. EM analysis of YSL CD63-pHluorin pDNA injected zebrafish revealed, qualitatively, that the former occurs, as we observed clusters of YSL derived exosomes labelled positive for aGFP immuno-gold, and mixed with dense aggregates in endo-lysosomal structures in endothelial cells of the caudal plexus (**Fig 5C**). This suggested a degradative fate of exosomes that we further investigated by treating zebrafish embryos with a specific inhibitor of the vacuolar H(+)-ATPase, Bafilomycin A1 (BafA), that elevates endosomal pH and decreases lysosomal degradation. Strikingly, incubation for 2h with 100nm BafA showed that endothelial cells of the vein plexus and the cardinal vein but not in the dorsal aorta endocytosed considerable amounts of CD63-pHluorin exosomes over time, mostly concentrating at peri-nuclear areas (**Fig 5D**, nuclei indicated with (N)). This accumulation, after only two hours of treatment, suggests that exosomes are mainly and continuously taken up by endothelial cells to be degraded in acidic endo-lysosomes.

We then tested whether exosomes could propagate beyond vascular endothelial walls. For this, we imaged the area of the caudal vein plexus and clearly observed labelled exosomes outside of the CVP (**Fig 5E, F; Suppl Movie 11**). Even though we observed some of the larger fluorescent exosome clusters concentrated adjacent to the CVP, we detected high levels of fluorescent exosomes outside of the vasculature throughout the whole embryo, notably visible where covering the muscle segments (**Suppl Movie 12**). The motility of the exosomes we observed outside the vasculature was irregular and resembled Brownian motion, clearly distinct from the dynamics of intracellular endosomal transport (**Suppl Movie 6**). However, in muscle cells, Bafilomycin A treatment (**Supp Video 13**) did not show accumulation of YSL derived exosomes as observed for endothelial cells. The large majority of signal was thus observed in endothelial cells and to a lesser extent in patrolling macrophages but not in other cell types, highlighting the specificity for targeted cells.

### Uptake of YSL-derived exosomes by endothelial cells relies on scavenger receptors and dynamin-dependent endocytosis

This specific interaction of labelled exosomes with target cell types likely relies on specific receptors (van Niel et al., 2018). Having this opportunity to visualize internalized CD63-pH exosomes with BafA treatment, we first proceeded to interfere with general uptake mechanism by inhibiting dynamin, a key regulator of clathrin dependent and independent endocytosis (Ferguson and De Camilli, 2012). We treated 2dpf YSL CD63-pHluorin expressing *Tg(kdrl:HRAS-mCherry)* embryos, expressing mCherry in the vasculature, with 3μM of dynamin inhibitor Pyrimidyn-7 (McGeachie et al., 2013) prior to BafA treatment. Strikingly, we observed a near complete block of exosome uptake by endothelial cells (**Fig 6A**). Mander’s coefficient (M2) of the green signal (exosomes) overlapping with the red signal (endothelium) revealed an average decrease of more than 10-fold of the uptake (**Fig 6B**). To exclude potential interference of random overlap in our analysis, we measured the Mander’s coefficient between the two channels with or without 180-degrees rotation of the green channel versus the red channel (**Fig 6C**). A diminished random overlap - comparable to the overlap observed in the Pyrimidyn-7 treated condition - showed that random colocalization does not significantly contribute to our measurements (**Fig 6C**). Similar quantification in ctrl or Pyrimidyn-7 condition showed a 5-fold reduction of uptake in macrophages (**Fig 6D**). We next focused on potential receptors able to specify this endocytosis to endothelial cells and macrophages.

**Figure 6.**
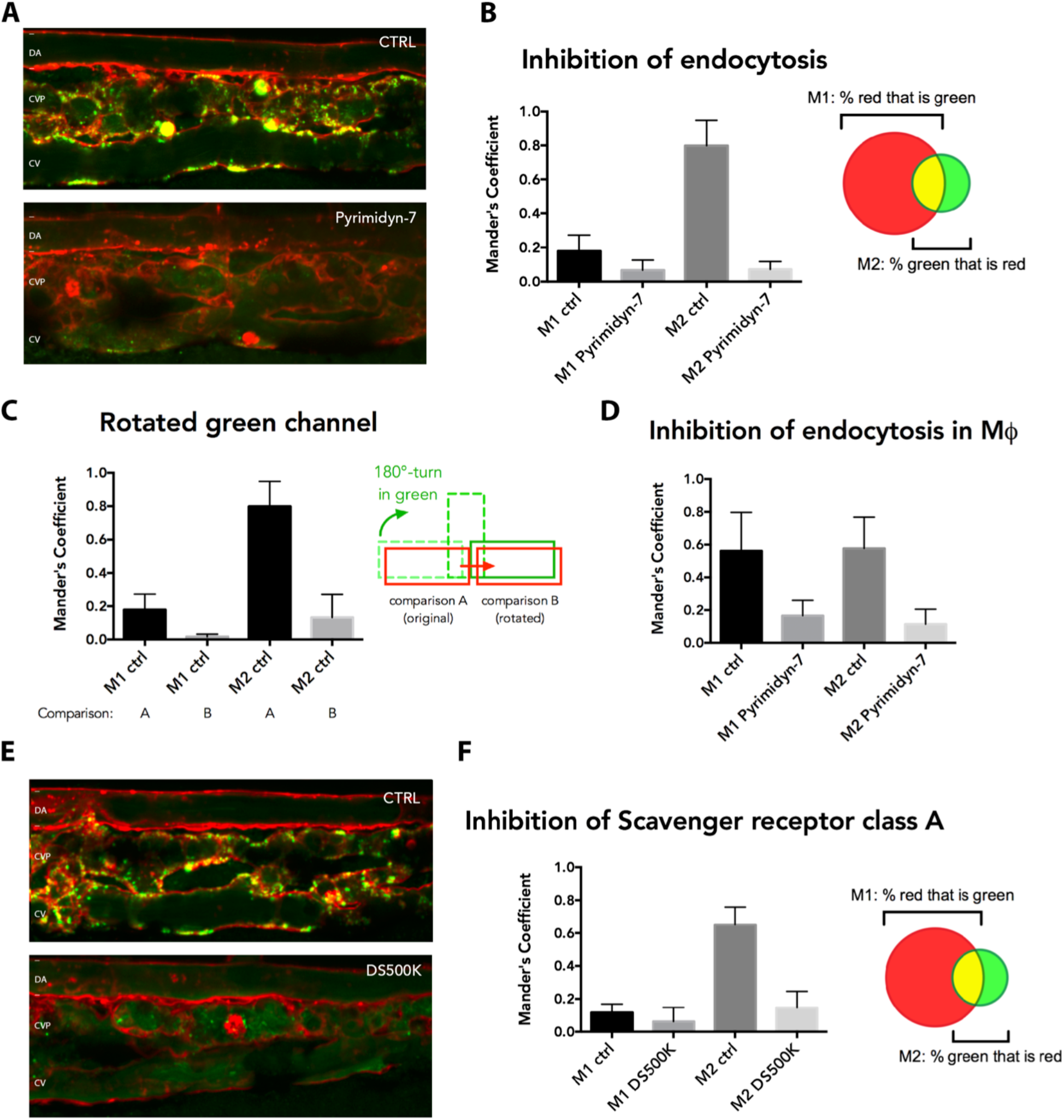
Uptake mechanism of YSL-EV in the endothelium. (A) Close-up of the CVP area of 3dpf *Tg(kdrl:Hsa.HRAS-mCherry)* zebrafish embryos expressing CD63-pHluorin in the YSL, control treated (DMSO) or treated with dynamin-inhibitor Pyrimidyn-7. BafA was used to show internalized EVs. (B) Quantification of experiments as performed in (e), showing the overlap-coefficient of red with green (M1) and green with red (M2) signal. (C) As in b, but with 180-degree rotation of green channel to control for random overlap. (D) As in a, but for macrophages alone. (E) Close-up of the CVP area of 3dpf *Tg(kdrl:Hsa.HRAS-mCherry)* zebrafish embryos expressing CD63-pHluorin in the YSL, non-injected or injected with 500K Dextran Sulfate (DS500K). BafA was used to show internalized EVs. (F) Quantification of experiments as performed in (b), showing the overlap-coefficient of red with green (M1) and green with red (M2) signal.

Interestingly, the CVP area harbors scavenger endothelial cells (SECs) that interact in a scavenger-receptor mediated manner with liposomes of 100nm injected in the circulation (Campbell et al., 2018). Scavenger receptors have been also reported in vitro to mediate exosome uptake (Plebanek et al., 2015). In particular, the Class A of Scavenger Receptors (SR-As) is expressed on endothelial cells and macrophages and is conserved in teleost (Bowdish and Gordon, 2009; Poynter et al., 2015) and mediate dynamin-dependent endocytocis (Fukuda et al., 2011; Jang et al., 2014). To investigate whether scavenger receptors are implicated in exosome uptake in vivo, we injected dextran sulfate (DexSO4-500K), a known competitive ligand for SR-A (Basu et al., 1979; Lysko et al., 1999), intravenously in CD63-pHluorin YSL expressing *Tg(kdrl:HRAS-mCherry)* embryos prior to BafA treatment. Strikingly, we observed a decrease of more than 4-fold of the green signal (EVs) overlapping with the red signal (endothelium) (**Fig 5G-H**), indicating that endogenous exosomes competed with DexSO4-500K for their uptake by the same class of receptors. This set of data revealed that *in vivo*, endogenous exosomes are specifically internalized by specialized veinous endothelial cells in a dynamin- and scavenger receptor dependent pathway to be targeted to lysosomes for degradation.

### Production of YSL-derived exosomes relies on a syntenin-dependent biogenesis pathway and support vasculogenesis of the caudal vein plexus

The enrichment for a Syntenin–Alix mediated machinery in our exosome (**Fig 3F**) (Baietti et al., 2012) led us to interfere with their generation and further investigate their potential function in vivo. A previous study in zebrafish reported the expression of syntenin-a, notably in YSL, at different stages of development (Lambaerts et al., 2012). Depletion of syntenin-a in the whole embryo in the same study revealed defects in epiboly progression and body-axis formation during early zebrafish development whereas depleting syntenin-a solely in the YSL only affected epiboly in 30% of the embryos and resulted in a well-formed axis in all embryos. We co-injected one of these validated morpholino oligonucleotides targeting the translation start site in syntenin-a transcripts (MOSyntA) or control morpholino (Ctrl MO) (Lambaerts et al., 2012) together with CD63-pHluorin pDNA specifically in the YSL of *Tg(kdrl:HRAS-mCherry)* embryos 4hpf. Of note, in our hands most embryos displayed normal development but showed a small delay in hatching and increase in mortality (**Fig 7A-C**). When considering only embryos with normal development (>50% in both conditions), whole body analysis showed a preserved expression CD63-pHluorin in the YSL in MOSyntA condition (**Fig 7A**) but a profoundly diminished number of fluorescent exosomes in the blood flow when compared to Ctrl MO condition (**Fig 7D,E**). This suggests that Syntenin is implicated in CD63 positive exosome biogenesis in the YSL *in vivo* and that endogenous exosome biogenesis can be manipulated. Incubation of these embryos for 2h with 100nm BafA confirmed the near absence of exosomes in the CVP, corresponding with a 20-fold reduction of the uptake of exosomes by endothelial cells in this region (**Fig 7F**). While we did not observe any major defect in angiogenesis or growth of MOSyntA embryos at 3dpf (**Fig 7C**), we did observe a decrease of ~18% in the height of the CVP between MOSyntA and MOCtrl treated embryos (**Fig 7G**), suggesting a role of exosomes produced from the YSL in the development of the CVP by taking them up and targeting them to lysosomes. Collectively, these data demonstrate that endogenous biogenesis of exosomes can be modulated and be used to reveal their potential physiological role in target cells.

**Figure 7.**
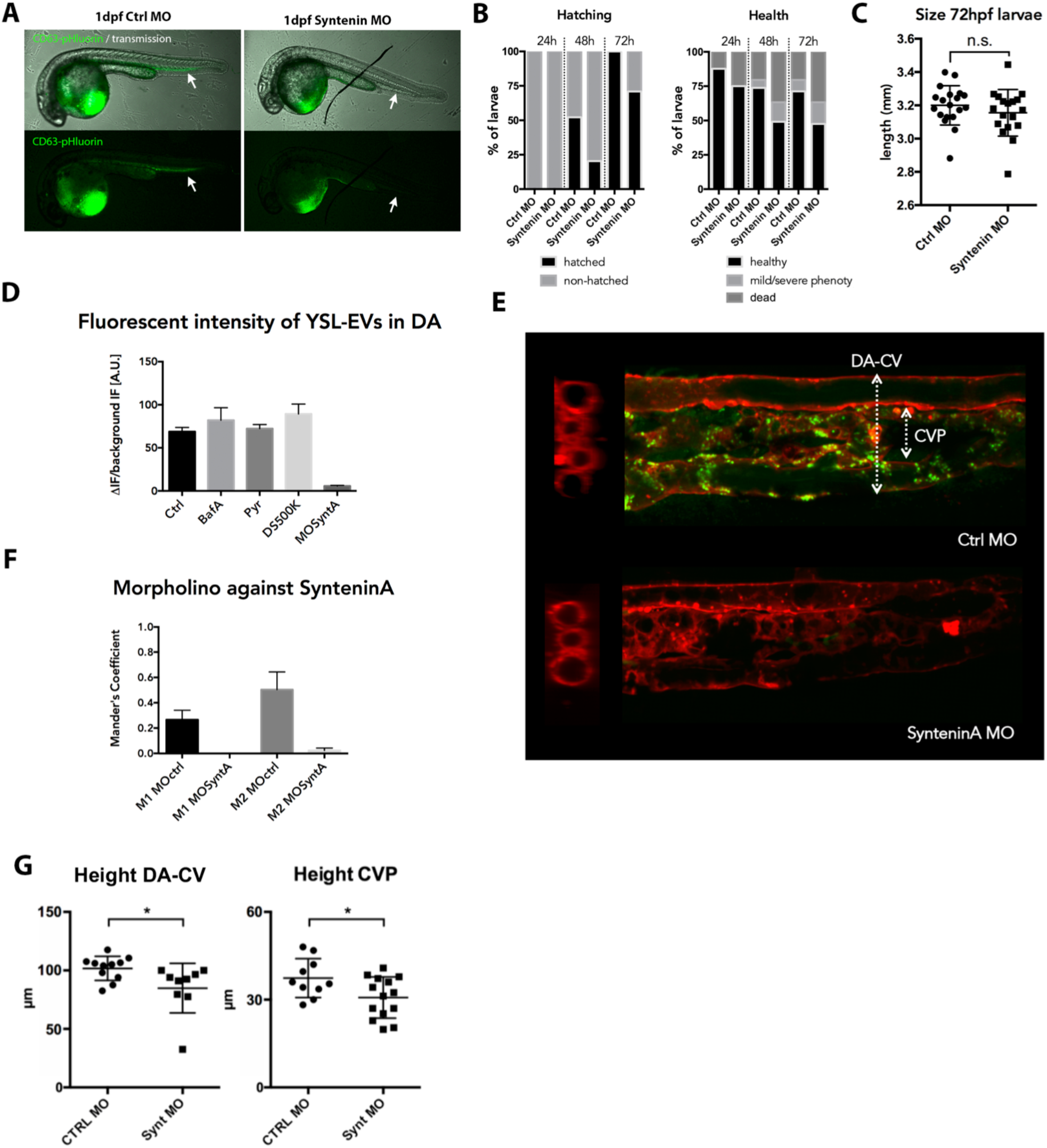
Functional analysis of YSL-EVs. (A) Transmission and fluorescent images of Ctrl MO or Syntenin MO treated 1dpf zebrafish embryos. (B) Quantification of different conditions of Ctrl MO or Syntenin MO treated 1-3dpf (24h-72h) zebrafish embryos. To the left, hatched vs non-hatched comparison. To the right, health status comparison with normal morphology (healthy), mild or severe developmental defects and mortality. (C) Size (length) of Ctrl MO or Syntenin MO treated 3dpf zebrafish embryos. (D) Fluorescent intensity of GFP signal in dorsal artery (DA) measured during different treatments. BafA=Bafilomycin-A, Pyr=Pyrimidyn-7, DS500K= 500KDa Dextran Sulfate, MOSyntA=Syntenin-A Morpholino, A.U.=arbitrary units. (E) Close-up of the CVP area of 3dpf *Tg(kdrl:Hsa.HRAS-mCherry)* zebrafish embryos expressing CD63-pHluorin in the YSL, co-injected with with MOctrl or MOSyntA in the YSL. DA=dorsal artery, CV=cardinal vein. (F) Quantification of overlap-coefficient of red with green (M1) and green with red (M2) in MOctrl or MOSyntA treated 3dpf zebrafish embryos. (G) Quantification of height of caudal part of vasculature, measured from DA to CV or specifically for the CVP at second intersegmental vessel (ISV).

## DISCUSSION

Here, we set up an in vivo model in zebrafish embryos to study various aspects of the physiology of endogenous exosomes in vivo at high spatio-temporal resolution. Our data reveal the release of exosomes from Danio rerio cell lines and embryos. Notably, our data highlight in vivo mechanisms for biogenesis, trafficking and uptake of endogenous exosomes using a combination of microscopic approaches. By proteomic analysis on purified YSL-exosomes, we could analyse their composition and unravel their nature and part of their biogenesis mechanism, extending to in vivo condition our fundamental knowledge of the molecular mechanisms regulating exosome biogenesis in vitro (Baietti et al., 2012). In the vasculature, we identified endothelial cells and macrophages of the Caudal Vein Plexus (CVP) as target cell types, corroborating the findings in the co-submitted study by Hyenne et al. Using this model, we were able to identify the fate of exosomes in target cells by showing a key role for dynamin-dependent endocytic process, which involved the class A Scavenger Receptor family (SR-A) in endothelial cells. Our data and the data of Hyenne et al also showed that, once endocytosed, a common and major fate of EVs and exosomes is lysosomal degradation, likely providing a trophic support to target cells. Finally, we could efficiently modulate exosome biogenesis in vivo and unveil their potential stimulatory role in vascular growth of the CVP. All in all, our data reveals the relevance of our model to investigate the origin, composition, biogenesis pathways, targets, fate and function of endogenous exosomes and EVs in vivo.

Until now, the development of a model to explore the release, journey and targets of endogenous exosomes *in vivo* had been hampered by the small size of the exosomes and the lack of appropriate methods to label them. A recent study reported the use of CD63-GFP in transgenic rats (Yoshimura et al., 2016). While this model amply demonstrates its usefulness for the *ex vivo* analysis of EV transfer to organs and bodily fluids and is relatively close to human, it is somewhat limited in its suitability for *in vivo* imaging. Here, we sought to fill that void by exploring a vertebrate model suitable for *in toto* imaging while maintaining sub-cellular and even supra-optical resolution. The CD63-pHluorin strategy has been recently shown to be an accurate in vitro model to explore new facets of the molecular mechanisms that control the biogenesis and secretion of exosomes (Verweij et al., 2018). Our study demonstrates that the CD63-pHluorin reporter could be used as previously shown in vitro (Verweij et al., 2018) to investigate the molecular regulation of exosome secretion. Such investigation is, however, still limited by a technological latch that hinders quantitative observation of bursts of fluorescence in cells that are not flat. We observed a relatively limited number of fusion events compared to transformed cell lines *in vitro*, in line with *in vitro* experiments indicating lower fusion activity with non-transformed cells (Verweij et al., 2018) and the increased presence of exosomes in bodily fluids of cancer patients (Baran et al., 2010; Galindo-Hernandez et al., 2013; Logozzi et al., 2009). The transposition of this tool in vivo also provided a model allowing single vesicle tracking from their site of production to their final destination using a combination of microscopic approaches. Our data show that zebrafish expressing CD63-pHluorin are a suitable *in vivo* model for the integrative study of individual EVs on tissue and sub-cellular level and are at the same time compatible with electron microscopic studies to reach supra-optical resolution. Live-tracking of endogenous exosomes in the whole organism of zebrafish embryos and notably their bodily fluids, opens new avenues to use this model to investigate the dynamics of body fluid flows with high precision (Bachy et al., 2008; Böhm et al., 2016), as observed in the cerebro-spinal fluid of CD63-pHluorin expressing zebrafish (**Sup Movie 12**).

We observed a large number of fluorescent vesicles after ubiquituous expression of CD63-pHluorin. Using tissue-specific expression, we could trace back the origin of these exosomes, and identified the Yolk Syncytial Layer (YSL), a secretory cell layer in direct contact with the blood as provider of endogenous exosomes for the rest of the organism. Mass spectrometry analysis of EVs derived from cell lines (AB.9, Zmel) and embryos in our study and in (Hyenne et al., co-submitted) showed a common composition that highlights the consistent presence of proteins involved in exosomes biogenesis by the syntenin-Alix-ESCRT-III pathway previously reported in vitro (Baietti et al., 2012). Of note, treatment of embryos with GW4869 did not affect exosome release from the YSL (**Sup Fig 1G**), suggesting that at least for this source of exosomes, ceramide is not a major regulator of their biogenesis (Trajkovic et al., 2008). These data suggest that in vivo, the syntenin-Alix-ESCRT-III pathway would be preponderant for exosomes biogenesis, in line with our morpholino experiments, opening new avenues to interfere with EVs generation and function in vivo. By defining a unique source of fluorescent exosomes by cell type specific expression, this tool is a relevant reporter to track exosome (sub)populations from their site of production to their final destination *in vivo*.

Our initial explorations uncovered endothelial cells and patrolling macrophages as recipient cells, strengthening the notion of targeted communication by exosomes. Although we cannot exclude a specific role of YSL derived exosomes once taken up by macrophages, we would favor the hypothesis that these scavenger cells may harvest EVs dwelling at the surface of endothelial cells to regulate the steady state level of EVs in the vasculature. In zebrafish, we show that endogenous exosomes are mainly targeted to the veinous endothelial cells of the CV(P). These cells show functional homology with specialized Liver Sinusoidal Endothelial Cells (LSECs) in higher vertebrates (Campbell et al., 2018) indicating interspecies correlation with the targeting of injected EVs to the liver in murine models (Charoenviriyakul et al., 2017; Di Rocco et al., 2016; Wiklander et al., 2015). LSECs clear soluble macromolecules and small particles from the circulation and in humans possess the highest endocytic capacity of all cells (Poisson et al., 2017). In older vertebrates, jawless, cartilaginous and bony fish, cells resembling the LSEC are located at different locations outside the liver and are collectively called Scavenger Endothelial Cells (SECs) that are similarly central to the removal and catabolism of a variety of macromolecules from the circulation (Seternes et al., 2002). Our results would place these YSL-derived exosomes alongside macromolecules. The lysosomal fate of YSL-derived exosomes in endothelial cells also echoes data showing that uptake and catabolism of exosomes in lysosomal compartments is likely a major fate as described in vitro (Tian et al., 2010). This pathway would allow degradation of EV content and provide a source of amino acids, nucleic acids, lipids and ions to recipient cells. Although we cannot rule out a structural function of YSL derived EVs during development, e.g. in patterning or differentiation (McGough and Vincent, 2016), the effect we observed on CV and CVP development when interfering with EVs biogenesis also suggests that endothelial cells may uptake and process EVs in lysosomes to use them as a source of macromolecules for their own development. We show that the uptake mechanisms of EVs by endothelial cells in the CV(P) implicate Scavenger Receptor and dynamin-dependent endocytosis. Further investigations are required to specify the precise endocytosis pathway of exosomes. Scavenger Receptors, such as Stabilin-2, have been recently described as mediating the uptake of 100nm sized liposomes in zebrafish (Campbell et al., 2018) and are expressed strongly on veinous endothelial cells of the CV (Wong et al., 2009). In line with the notion of a trophic role of EVs by their degradation, various scavenger receptors internalize their ligands in a dynamin-dependent manner and traffic to lysosomes (Fukuda et al., 2011) morphants for one of them, CL-P1, show malformation of vascular vessels and subsequent developmental defects.

The presence of numerous EVs in the bloodstream but also in the ISF of developing zebrafish embryos strengthens ex-vivo observations of the presence of EVs in both fluids (Gromov et al., 2013; Johnstone et al., 1989). Recent evidence indicates that exosomes or EVs containing miRNAs can also in- and extravasate biological barriers to access in ISF under physiological conditions (Thomou et al., 2017) but the YSL in zebrafish is directly connected with the ISF, providing a direct way for exosomes from YSL to access the ISF. Interstitial fluid contains amino acids, sugars, fatty acids, solutes, and other cellular products and provides support for tissue cells and removes metabolic waste (Renkin and Crone, 1996). The motility of the CD63-pHluorin positive exosomes we observed outside the vasculature was irregular and resembled Brownian motion, clearly distinct from the dynamics of intracellular endosomal transport (**Suppl Video 6**). Of note, Bafilomycin A treatment (as in **Fig 5D**) did not show accumulation of YSL derived exosomes in muscle cells (**Suppl Video 13**). Such observation would exclude uptake and lysosomal degradation as a way to use EVs as a source of macromolecules by other cells than endothelial cells. However, our proteomic analysis revealed the presence and enrichment for various lipid-, amino acid and ion transporters that would support a role of YSL-derived exosomes as ‘macromolecule dispenser’ to surrounding cells, as these transporters could potentially mediate the bioavailability of exosomal contents (Allen and Cullis, 2013). All in all, the presence of these EVs in various bodily fluids supports a general role in metabolism, e.g. in nutrient delivery, and makes a specific role in other YSL functions, e.g. embryonic patterning, less likely. Nevertheless, we cannot exclude that these EVs are used to exchange genetic information between cells in vivo as shown recently (Ashley et al., 2018; Pastuzyn et al., 2018; Zomer et al., 2015) and future development of adapted tools as described in vitro (Yim et al., 2016) are required to explore these functions in detail.

Despite its several advantages, we cannot exclude that in this model, expression of this reporter might affect exosome secretion or composition. Nevertheless, nanoparticle tracking did not show a significant change of the number of particles between non-injected and CD63-pHluorin YSL expressing embryos, and proteomics indicated the presence of endogenous zCD63 in the YSL-CD63-pHluorin labelled exosomes. Another potential downside of our approach might be that expression of a TSPAN-reporter may only label a subpopulation of exosomes as CD63 may not be equally enriched in all exosome subpopulation (Kowal et al., 2016). Isolating a subpopulation from the heterogenous family of EVs with this reporter would however present the advantage to target specific fate or functions. Future studies could benefit for this purpose from using zebrafish CD63-pHluorin and/or the expression of other TSPAN-pHluorin markers, including CD81, CD82, or CD9 (Verweij et al., 2018). Additionally, EVs that bud from the plasma membrane might also contain CD63-pHluorin, meaning that once secreted there is no straight-forward way of distinguishing exosomes from other EVs. Future studies using other tetraspanins such as CD9 or CD81 that are more enriched at the plasma membrane than CD63 could perhaps aid in this distinction. The present study was restricted to the YSL, but development of future strains with cell type specific expression will allow to investigate the various roles of exosomes secreted under physiological and pathological conditions by nearly all cell types investigated so far. In addition, combining cell-type specific expression of TSPAN-pHluorin markers with an IP for GFP and proteomic analysis of exosomes could potentially be used to discover pathological bio-markers for each producing cell type targeted in vivo.

Importantly, the study of Hyenne et al (co-submitted) also used zebrafish embryos to investigate the behaviour and fate of systemically injected exogenous exosomes originating from zebrafish melanoma cells. Whereas previous studies have indicated how tumour-exosome integrins determine their organotropism (Hoshino et al., 2015), the combination of both models presented here reveals common uptake mechanisms between exogenous/pathological exosomes and endogenous/physiological exosomes exist, an interesting phenotype that has never been explored before at this scale. How circulating EVs target specific cell types at distance remains to be solved, mostly because this step could not be captured before, but the *in vivo* model presented here opens new avenues to unveil such mechanisms under physiological and pathological conditions. The genetic and structural homologies between zebrafish and human open new perspectives to validate exosome-mediated intercellular communication in the complex architecture of the organism, notably inter-organ communication and their ability to cross biological barriers. The fast development of disease models in zebrafish also provides the opportunity to tackle putative roles of exosomes in several pathologies that involves complex network of cell types. Additionally, mosaic expression resulting from pDNA injection will allow stochastic labeling of cells within a tissue or throughout the organism, depending on the promotor associated to the CD63-pHluorin construct, facilitating single-cell imaging and single-EV tracking. We therefore propose the zebrafish embryo expressing fluorescent markers for exosomes (e.g. CD63-pHluorin) as a new model to study endogenous EVs *in vivo*, which will open new avenues to unravel fundamental aspects in EV physiology and assess their clinical applications. Altogether the study from Hyenne et al and ours showcase the variety of questions that could likely be investigated in model organisms to validate the numerous in vitro studies that reported on the biology and function of EVs so far (Yáñez-Mó et al., 2015).

## METHODS

### Reagents

Lidocaine (Sigma-Aldrich, L7757) was dissolved in ethanol and used at 640μM, Bafilomycin A (Sigma, SML1661) was used at 200nM, Dextran Sulfate (Sigma, D8906) was dissolved in H2O and used at 20mg/mL. BODIPY C12 558/568 (Life Technologies SAS, D3835) was injected in the yolk at 1.5 ng/nl BODIPY C12, using canola oil as lipid carrier. Collagenase I was used at 200mg/mL (ThermoFisher, 17018029). pUbi-CD63-pHluorin was constructed by first cloning CD63-pHluorin (human CD63; sequence identical to Verweij et al., 2018) using primer pairs 5′-GGGGACAAGTTTGTACAAAAAAGCAGGCTGGatggcggtggaaggagga-3′ and 5′-GGGGACCACTTTGTACAAGAAAGCTGGGTCctacatcacctcgtagccacttct-3′ into pDONR221 (Gateway, Invitrogen). pDONR221-CD63-pHluorin was subsequently recombined with p5E-ubi (Addgene plasmid # 27320), p3E-polyA 153, and pDEST R4-R3 (Gateway, Invitrogen) to create pUbi-CD63-pHluorin.

### Ethic statements

All fish are housed in the fish facility of our laboratory, which was built according to the local animal welfare standards. All animal procedures were performed in accordance with French and European Union animal welfare guidelines.

### Zebrafish strains

Zebrafish (*Danio rerio*, AB9, Casper, Tg(kdrl:Hsa.HRAS-mCherry), Tg(mpeg1:mCherryF) strains) were staged and cared for according to standard protocols (Westerfield).

### Injections

To induce expression of CD63-pHluorin, embryos were injected at the one-cell stage (for ubiquitous expression) or at 4h in the YSL (for YSL-specific expression). On the next day, injected embryos were inspected under a stereomicroscope. Only embryos that developed normally were assayed.

### Morpholino (MO)

The MO against zSyntenin-A (5′-TACAACGACATCCTTTCTGCTTTCA-3’) was purchased from GeneTools and was used at 1:4 dilution from a 1 mM stock.

### Live imaging

Embryos were anesthetized with tricaine and embedded in 1.5% low melting point agarose. For most experiments, embryos were imaged at 3dpf. Recordings were performed at 28°C using a Nikon TSi spinning-wide (Yokagawa CSU-W1) microscope.

### Image analysis

All imaging data was analysed using ImageJ. Mander’s overlap coefficient was determined using the JACoP plugin.

### Statistical analysis

We performed statistical analysis (Student’s two-tailed t-test for significance) using GraphPad Prism version 6.0 (GraphPad software). Unpaired analysis was used unless specified otherwise. Data distribution was assumed to be normal but this was not formally tested.

### Exosome preparation

Exosomes were prepared from the supernatant of 24h cultured HeLa cells as described before (Théry et al., 2006). Briefly, exosomes were purified from the cultured media with exosome-free serum. Differential centrifugations at 500 g (2 × 10 min), 2,000 g (2 × 15min), 10,000g (2 × 30min) eliminated cellular debris and centrifugation at 100,000 g (60 min) pelleted exosomes. The exosome pellet was washed once in a PBS, centrifuged at 100,000 g for 1 h and re-suspended in 50-200 μL PBS, depending on the starting volume.

### Label-free Mass-spectrometry

Exosomes were purified by differential ultracentrifugation as previously described (Théry et al., 2006; Verweij et al., 2013), and followed (for YSL derived exosomes) by an IP for GFP using GFP-trap. Pull-downs were performed in biological replicates (n = 3) To characterize the exosome proteins, we qualitatively and quantitatively analyzed them by Orbitrap Fusion Tribrid mass spectrometry, by using a label free approach. For exosome quantification, SDS/PAGE was used without separation as a clean-up step, and only one gel slice was excised otherwise proteins were separated on SDS/PAGE and four gel slices were excised. Subsequently, gel slices were washed and proteins were reduced with 10 mM DTT before alkylation with 55 mM iodoacetamide. After washing and shrinking the gel pieces with 100% MeCN, we performed in-gel digestion using trypsin/LysC (Promega) overnight in 25 mM NH4HCO3 at 30 °C. Peptide were then extracted using 60/35/5 MeCN/H2O/HCOOH and vacuum concentrated to dryness Samples were chromatographically separated using an RSLCnano system (Ultimate 3000, Thermo Scientific) coupled to an Orbitrap Fusion Tribrid mass spectrometer (Thermo Scientific).

Peptides were loaded onto a C18 column (75-μm inner diameter × 2 cm; nanoViper Acclaim PepMapTM 100 or 300 μm × 5 mm PepMapTM 100, 5 μm, Thermo Scientific), separated and MS data acquired using Xcalibur software. Peptides separation was performed over a linear gradient of 100 min from 5% to 25% or 5% to 35% MeCN (75-μm inner diameter × 50 cm; nanoViper C18, 2 μm or 5 μm, 100Å, Acclaim PepMapTM RSLC, Thermo Scientific). Full-scan MS was acquired in the Orbitrap analyzer with a resolution set to 120,000 and ions from each full scan were HCD fragmented and analyzed in the linear ion trap.

For identification the data were searched against the UniProtKB Zebrafish database using Sequest through proteome discoverer. Enzyme specificity was set to trypsin and a maximum of two missed cleavage site were allowed. Oxidized methionine, N-terminal acetylation, and carbamidomethyl cysteine were set as variable modifications. Maximum allowed mass deviation was set to 10 ppm for monoisotopic precursor ions and 0.6 Da for MS/MS peaks. The resulting files were further processed using myProMS (Poullet et al., 2007). FDR calculation used Percolator and was set to 1%. The label free quantification was performed by peptide Extracted Ion Chromatograms (XICs) computed with MassChroQ version 1.2.1 (Valot et al., 2011). Global MAD normalization was applied on the total signal to correct the XICs for each biological replicate. Protein ratios were computed as the geometrical mean of related peptides. To estimate ratio significance, a two-tailed t test was performed and p-values were adjusted with a Benjamini–Hochberg FDR procedure with a control threshold set to 0.05. Fold change-based GO enrichment analysis was performed as in Kowal et al. (Kowal et al., 2016).

## SUPPLEMENTARY MATERIAL LEGENDS

**Sup Figure 1.**
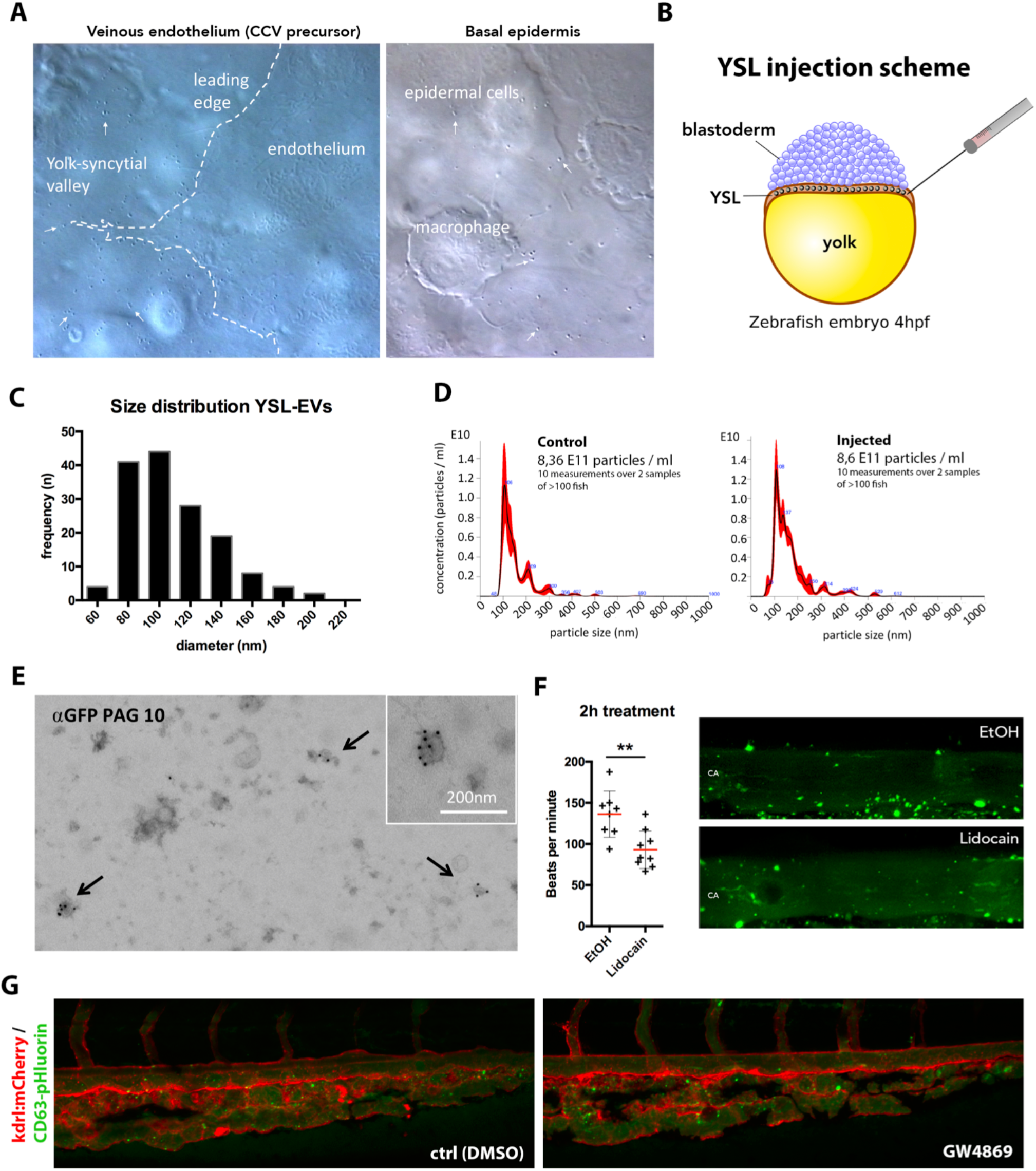
(A) Video-enhanced Nomarski microscopy of the large interstitial space between the YSL and the overlying epidermis around the onset of blood circulation (~24hpf) of a zebrafish embryo, showing numerous vesicles. White arrows indicate extracellular vesicles. (B) Schematic of YSL injection at 1K cell stage. (C) Size-distribution of 150 vesicles on the YSL (fig 3c). (D) NTA-analysis of particle concentration of CD63-pHluorin YSL injected (left) and non-injected fish. (E) EM analysis on vesicles isolated from dissociated YSL CD63-pHluorin expressing 3dpf zebrafish embryos, labelled against GFP with PAG10. (F) Effect of blood flow manipulation on attachment of EVs to the CA. To the left: graph showing effect of lidocaine on heart-beat (pacemaker activity). To the right: representative example of EV accumulation in treated (lido) condition and ctrl. (G) Overview of the CVP area of a 3dpf Tg(kdrl:Hsa.HRAS-mCherry) zebrafish embryo expressing CD63-pHluorin in the YSL, incubated with GW4869 or ctrl treated.

**Sup Video 1** | Real-time video of blood vessel posterior to the head of a 3dpf CD63-pHluorin ubiquitously expressing zebrafish embryo.

**Sup Video 2** | Real-time video of blood vessel in the gills of a 3dpf CD63-pHluorin ubiquitously expressing zebrafish embryo.

**Sup Video 3** | Real-time video of blood vessel in the brain of a 3dpf CD63-pHluorin ubiquitously expressing zebrafish embryo.

**Sup Video 4** | Real-time video of the cardinal vein of a 3dpf CD63-pHluorin YSL expressing zebrafish embryo.

**Sup Video 5** | Time-lapse of the CVP area of a 3dpf Tg(kdrl:Hsa.HRAS-mCherry) zebrafish embryo expressing CD63-pHluorin in the YSL.

**Sup Video 6** | CVP area imaged at high-speed in a zebrafish embryo expressing CD63-pHluorin in the YSL (combination of three consecutively acquired stacks).

**Sup Video 7** | Close-up of CVP and dorsal vein area in Tg(mpeg1:mCherryF) zebrafish embryo expressing CD63-pHluorin in the YSL.

**Sup Video 8** | Close-up of a macrophage in the CVP area of a 3dpf Tg(kdrl:Hsa.HRAS-mCherry) zebrafish embryo expressing CD63-pHluorin in the YSL.

**Sup Video 9** | Time-lapse of a macrophage in the CVP area of a 3dpf BODIPY-C12 injected zebrafish embryo expressing CD63-pHluorin in the YSL.

**Sup Video 10** | Real-time video of the cardinal vein of a 3dpf CD63-pHluorin YSL expressing zebrafish embryo.

**Sup Video 11** | 3D-reconstruction of the mid-trunk segment of of 3dpf zebrafish embryos expressing CD63-pHluorin in the YSL and mCherry under the ubiquitous RPL5 promoter.

**Sup Video 12** | Time-lapse of the neural tube of 3dpf CD63-pHluorin YSL expressing zebrafish. Video play-back speed is half

**Sup Video 13** | 3D-reconstruction of the mid-trunk segment of of 3dpf zebrafish embryos expressing CD63-pHluorin in the YSL and mCherry under the ubiquitous RPL5 promoter and treated with 200nM BafA for 2h.

**Sup Table I** | Proteomic analysis on exosomes isolated from AB.9 caudal fibroblast cells

**Sup Table II** | Proteomics analysis on ex vivo isolated EVs from 3dpf zebrafish embryos – enrichment of proteins form YSL-derived EVs vs all EVs

## ACKNOWLEDGEMENTS

We acknowledge Vincent Fraisier, Ludovic Leconte and Christine Viaris for technical assistance and Pascale Zimmermann for stimulating discussions. The authors greatly acknowledge the Cell and Tissue Imaging (PICT-IBiSA) and Nikon Imaging Centre, Institut Curie, member of the French National Research Infrastructure France-BioImaging (ANR10-INBS-04)”.

Financing support from Fondation pour la Recherche Medicale (contract AJE20160635884),) to G.v.N., the European Molecular Biology Organization grant (EMBO ALTF 1383-2014),) to F.V., the Fondation ARC pour la Recherché sur le Cancer fellowship (PJA 20161204808),) to F.V., LabEx CeTtisPhyBio to G.v.N. and F.V., and a “Région Ile-de-France” grant to D.L.

